# Meta-analysis of targeted metabolomics data from heterogeneous biological samples provides insights into metabolite dynamics

**DOI:** 10.1101/509372

**Authors:** Ho-Joon Lee, Daniel M. Kremer, Peter Sajjakulnukit, Li Zhang, Costas A. Lyssiotis

## Abstract

**Introduction:** Mass spectrometry-based metabolomics coupled to liquid chromatography, or LC-MS metabolomics, has become the most popular tool for global metabolite abundance profiling to study metabolism. However, the physicochemical complexity of metabolites poses a major challenge for reliable measurements of metabolite abundance. One way to address the issue is to use multiple chromatographic methods to capture a greater range of molecular diversity. We previously developed a tandem mass spectrometry-based label-free targeted metabolomics analysis framework coupled to two distinct chromatographic methods, reversed-phase liquid chromatography (RPLC) and hydrophilic interaction liquid chromatography (HILIC), with dynamic multiple reaction monitoring (dMRM) for simultaneous detection of over 200 metabolites to study core metabolic pathways.

**Objectives:** We aim to analyze a large-scale heterogeneous data compendium generated from our LC-MS/MS platform with both RPLC and HILIC methods to systematically assess measurement quality in biological replicate groups and to gain insights into metabolite dynamics across different biological conditions.

**Methods:** Our metabolomics framework was applied in a wide range of experimental systems including cancer cell lines, tumors, extracellular media, primary cells, immune cells, organoids, organs (e.g. pancreata), tissues, and sera from human and mice. We also developed computational and statistical analysis pipelines, which include hierarchical clustering, replicate-group CV analysis, correlation analysis, and case-control paired analysis.

**Results:** We generated a compendium of 42 heterogeneous deidentified datasets with 635 samples using both RPLC and HILIC methods. There exist signature metabolites that correspond to heterogeneous phenotypes, involved in several metabolic pathways. The RPLC method shows overall better reproducibility than the HILIC method for most metabolites including polar amino acids. Correlation analysis reveals high confidence metabolites irrespective of experimental systems such as methionine, phenylalanine, and taurine. We also identify homocystine, reduced glutathione, and phosphoenolpyruvic acid as highly dynamic metabolites across all case-control paired samples.

**Conclusions:** Our study is expected to serve as a resource and a reference point for a systematic analysis of label-free LC-MS/MS targeted metabolomics data in both RPLC and HILIC methods with dMRM.

## 1. Introduction

Cellular metabolism encompasses all the reactions inside a cell that involve small molecules. In order to understand metabolism and its regulation on a global level, metabolomics approaches have become essential for detecting metabolites and measuring their concentrations. Technological advancement of mass spectrometry (MS) has made it a popular and powerful platform for metabolomics studies (Johnson et al., 2016; Patti et al., 2012; Wishart, 2016). Although nuclear magnetic resonance (NMR)-based metabolomics is also widely used, MS is more easily coupled to various chromatographic columns to separate analytes prior to analysis, thereby reducing the complexity of a biological sample and increasing sensitivity for simultaneous detection of a large number of metabolites (Griffiths et al., 2010; Zhou and Yin, 2016).

Metabolomics can be performed as either targeted or non-targeted. Targeted metabolomics aims to detect and quantify a pre-defined list of specific metabolites, whereas non-targeted or untargeted metabolomics aims to detect and quantify all known and unknown metabolites or compounds in a sample as an unbiased discovery strategy (Dudley et al., 2010; Fuhrer and Zamboni, 2015). Untargeted metabolomics has more challenges than targeted metabolomics, both technically and computationally, and hence its use is more limited than targeted approaches, even in the discovery arena. For example, targeted metabolomics has been used routinely to study well-known metabolic pathways, such as glycolysis, the tricarboxylic acid (TCA) cycle, and nucleotide and amino acid biosynthesis pathways. Nevertheless, detection and quantification of specific metabolites in these pathways can still be challenging, particularly if one aims to target hundreds of metabolites in a single experiment.

MS-based metabolomics is performed predominantly by subjecting samples to chromatographic separation before MS analysis. Chromatographic columns make it possible to separate complex analyte mixtures based on physicochemical properties of a wide range of compounds, including isomers. Liquid chromatography (LC) is most frequently used, while gas chromatography (GC) is preferred for measuring volatile compounds. LC methods include reversed-phase liquid chromatography (RPLC) and hydrophilic interaction liquid chromatography (HILIC). RPLC is typically used for a broad range of metabolites, especially nonpolar and weakly polar metabolites, whereas HILIC has a complementary usage for hydrophilic, polar, and ionic metabolites such as sugars, amino acids, and nucleic acids (Buszewski and Noga, 2012; Cubbon et al., 2010; Ivanisevic et al., 2013; Lu et al., 2008; Rojo et al., 2012; Tang et al., 2016). While a recent study performed a systematic technical evaluation of hydrophilic and hydrophobic LC-MS in metabolomics profiling (Xie et al., 2018), it lacks a biological evaluation of abundance measurements in biological replicates under a wide range of different conditions.

We previously developed a tandem mass spectrometry-based targeted metabolomics system that profiles abundance of more than 200 metabolites as a steady-state snapshot of global metabolism. It utilizes both RPLC and HILIC methods with dynamic MRM (dMRM) as a means to maximize the coverage and sensitivity of target metabolites. Together with our customized computational and statistical analysis pipeline, this platform has been recently applied in several biological contexts (Halbrook et al., 2018; Schofield et al., 2018; Sousa et al., 2016; Svoboda et al., 2018). Herein, we report heterogeneous data sets from a wide range of experiments and sample types that were generated on our LC-MS/MS targeted metabolomics platform over a period of one year. Using these, we carried out comprehensive global meta-analysis in which we systematically evaluated data quality of both RPLC and HILIC methods by statistical measures and characterized global patterns of metabolite changes and variability in case versus control groups.

## 2. Materials and methods

### 2.1. Sample preparation

Preparation of the sample types below has been described before (Halbrook et al., 2018; Schofield et al., 2018; Sousa et al., 2016; Svoboda et al., 2018; Yuan et al., 2012). A brief description follows.

#### Primary and cultured cells

Biological replicates with n ≥ 3 were seeded at equivalent density, grow in log phase, and at end point media was fully aspirated. Aqueous metabolites of adherent primary or cultured cells on 6-well or 10cm^2^ plates were extracted by adding 1 mL or 4 mL of 80% cold (−80°C) methanol, respectively, followed by incubation at −80°C for 10 min. Cells were then scrapped and all material, including soluble and insoluble material, was collected. This was then followed by centrifugation at 14,000 rpm at 4°C for 10 min to pellet the insoluble material. Suspension cells were gently centrifuged to pellet and media was completely aspirated. Metabolites were extracted using the method above. The procedure is done on a bucket of dry ice and as quickly as possible in order to stop metabolism immediately. Samples were normalized by the volume corresponding to protein concentration measured from parallelly prepared lysates (typically, 1 – 2 million cells at 70 – 80% confluence). Then, samples were dried under vacuum and suspended in a 1:1 H2O/methanol solution for LC-MS analysis.

#### Tissues, organs, and tumors

Samples of 50 – 200 mg (n ≥ 3) were placed in a tube containing 1 mL of 80% cold (−80°C) methanol and then homogenized using steel beads and a Qiagen Tissue Lyser by multiple rounds of 45-second shaking at room temperature before centrifugation at 14,000 rpm at 4°C for 10 min. Samples were normalized by taking the volume corresponding to 10 mg of the tumor weight and further processed as above for LC-MS analysis.

#### Cultured media and serum

Metabolites from equivalent volumes of media or serum (n ≥ 3; typically, ~200 μl) were extracted by adding 100% cold (−80°C) methanol with a 1:4 ratio of the sample to methanol (i.e., 80% methanol final) and further processed as above for LC-MS analysis.

### 2.2. LC-MS metabolomics analysis

Our LC-MS/MS metabolomics analysis was performed as described previously (Halbrook et al., 2018; Schofield et al., 2018; Sousa et al., 2016). In brief, an Agilent 1290 UHPLC and 6490 Triple Quadrupole (QqQ) Mass Spectrometer (LC-MS) were used for label-free targeted metabolomics analysis. Agilent MassHunter Optimizer and Workstation Software LC-MS Data Acquisition for 6400 Series Triple Quadrupole B.08.00 was used for standard optimization and data acquisition. Agilent MassHunter Workstation Software Quantitative Analysis Version B.0700 for QqQ was used for initial raw data analysis.

For RPLC, a Waters Acquity UPLC BEH TSS C18 column (2.1 × 100mm, 1.7μm) was used in the positive ionization mode with mobile phase (A) consisting of 0.5 mM NH_4_F and 0.1% formic acid in water; mobile phase (B) consisting of 0.1% formic acid in acetonitrile. Gradient program: mobile phase (B) was held at 1% for 1.5 min, increased to 80% at 15 min, then to 99% at 17 min and held for 2 min before going to initial condition and held for 10 min. For HILIC, a Waters Acquity UPLC BEH amide column (2.1 × 100mm, 1.7μm) was used in the negative ionization mode with mobile phase (A) consisting of 20 mM ammonium acetate (NH_4_OAc) in water at pH 9.6; mobile phase (B) consisting of acetonitrile (ACN). Gradient program: mobile phase (B) was held at 85% for 1 min, decreased to 65% at 12 min, then to 40% at 15 min and held for 5 min before going to the initial condition and held for 10 min.

Both columns were at 40 ̊C and 3 μl of each sample was injected into the LC-MS with a flow rate of 0.2 ml/min. Calibration was achieved through Agilent ESI-Low Concentration Tuning Mix. Optimization was performed on the 6490 QqQ in the RPLC-positive or HILIC-negative mode for each of 245 standard compounds (215 and 217 compounds for RPLC-positive and HILIC-negative, respectively) to obtain the best fragment ion and MS parameters such as fragmentation energy for each standard. Retention time (RT) for each standard was measured from a pure standard solution or a mix standard solution. The LC-MS/MS methods were created with dynamic MRM (dMRM) with RTs, RT windows, and transitions of all 245 standard compounds (see Supplementary Table). Key parameters of electrospray ionization (ESI) in both the positive and the negative acquisition modes are: Gas temp 275°C, Gas Flow 14 l/min, Nebulizer at 20 psi, SheathGasHeater 250°C, SheathGasFlow 11 l/min, and Capillary 3000 V. For MS: Delta EMV 200V or 350V for the positive or negative acquisition mode respectively and Cycle Time 500ms and Cell Acc 4V for both modes. In this study we denote the dMRM method with RPLC in the positive ionization mode by RPLC-Pos-dMRM and the dMRM method with HILIC in the negative ionization mode by HILIC-Neg-dMRM. We note that our methods do not distinguish stereoisomers, hence care should be taken in interpretation of such data.

### 2.3. Computational data processing, quality control, and statistical analysis

Pre-processed data with Agilent MassHunter Workstation Software Quantitative Analysis were post-processed for further quality control in the programming language R. First, we examined the distribution of sums of all metabolite abundance peak areas across individual samples in a given experiment as a measure for equal sample loading into the instrument. Any outlier sample was removed, which is defined by a loading difference of greater than 70% compared to the average of the total abundance sums. Next, we calculated coefficients of variation (CVs) in all groups of biological replicate samples (n ≥ 3) for each metabolite given a cut-off value of peak areas in each of the RPLC-Pos-dMRM and the HILIC-Neg-dMRM methods. We then compared distributions of CVs for the whole dataset for a set of peak area cut-off values of 0, 1000, 5000, 10000, 15000, 20000, 25000 and 30000 in each method. A noise cut-off value of peak areas in each method was chosen by manual inspection of the CV distributions. The noise-filtered data of individual samples were then normalized by the total intensity of all metabolites. We retained only those metabolites with at least 2 replicate measurements for a given experimental variable. The remaining missing value in each condition for each metabolite was filled with the median value of the other replicate measurements. Then, each metabolite abundance level in each sample was divided (i.e., scaled) by the mean of all abundance levels across all samples in a given experiment for comparisons, statistical analyses, and visualizations among metabolites. Finally, we visually inspected a correlation heatmap profile of the samples of the resultant data to identify and remove any further outlier samples based on hierarchical clustering and abnormal heatmap patterns. Heatmaps were generated with the function, *heatmap.2*, from the Bioconductor package, *gplots*, and hierarchical clustering was performed with default parameters in *heatmap.2*. Pathway analysis was done using the webtool, MetaboAnalyst (Chong et al., 2018).

## 3. Results and discussion

### 3.1. Global view of a compendium of heterogeneous targeted metabolomics datasets

We compiled 42 datasets of LC-MS/MS-based targeted metabolomics with both RPLC-Pos-dMRM and HILIC-Neg-dMRM methods. There are a total of 638 samples analyzed, which include cancer cell lines, tumors, extracellular media, primary cells, immune cells, organoids, organs (e.g. pancreata), tissues, sera, and shRNA/drug treatments in human and mice. A total of 448 measurements were made in both methods for 285 unique compound entities (due to multiple detection). We retained measurements for those metabolites detected in both RPLC-Pos-dMRM and HILIC-Neg-dMRM methods as independent measurements. From this compendium we constructed a 448 by 638 data matrix for global characterization. 36.7% of the values in this data matrix were missing, and we removed 11 samples where no compound was measured. Figure 1A depicts a correlation heatmap and a hierarchical clustering of all metabolites in both RPLC-Pos-dMRM and HILIC-Neg-dMRM methods. Figure 1B shows a correlation heatmap and a hierarchical clustering of all samples. We find that there are a few groups of highly correlated metabolites. For example, at the height of 6 in the dendrogram tree, the 31 metabolites in Cluster #1 are enriched in purine, pyrimidine, glycine/serine, and glutathione metabolism, and the 88 metabolites in Cluster #2 include metabolites in the TCA cycle, glycolysis, pyruvate, and phenylalanine/tyrosine/tryptophan metabolism (Figure 1C). This suggests that those metabolites tend to change their abundance in the same direction across different conditions, which suggests that they are subject to pathway level metabolic regulation. The lack of clustering among samples reflects the heterogeneous nature of all datasets and indicates that experimental bias is not driving clustering and influencing downstream analysis. In fact, the most highly correlated 25 samples (Cluster #3, Figure 1B) do not exhibit common phenotypes, and this cluster includes cell lines, murine tumors and several experimentally distinct variables. Therefore, it suggests the existence of general metabolic phenotypes that can arise from various perturbations, independent of cell/tissue type.

**Figure 1.**
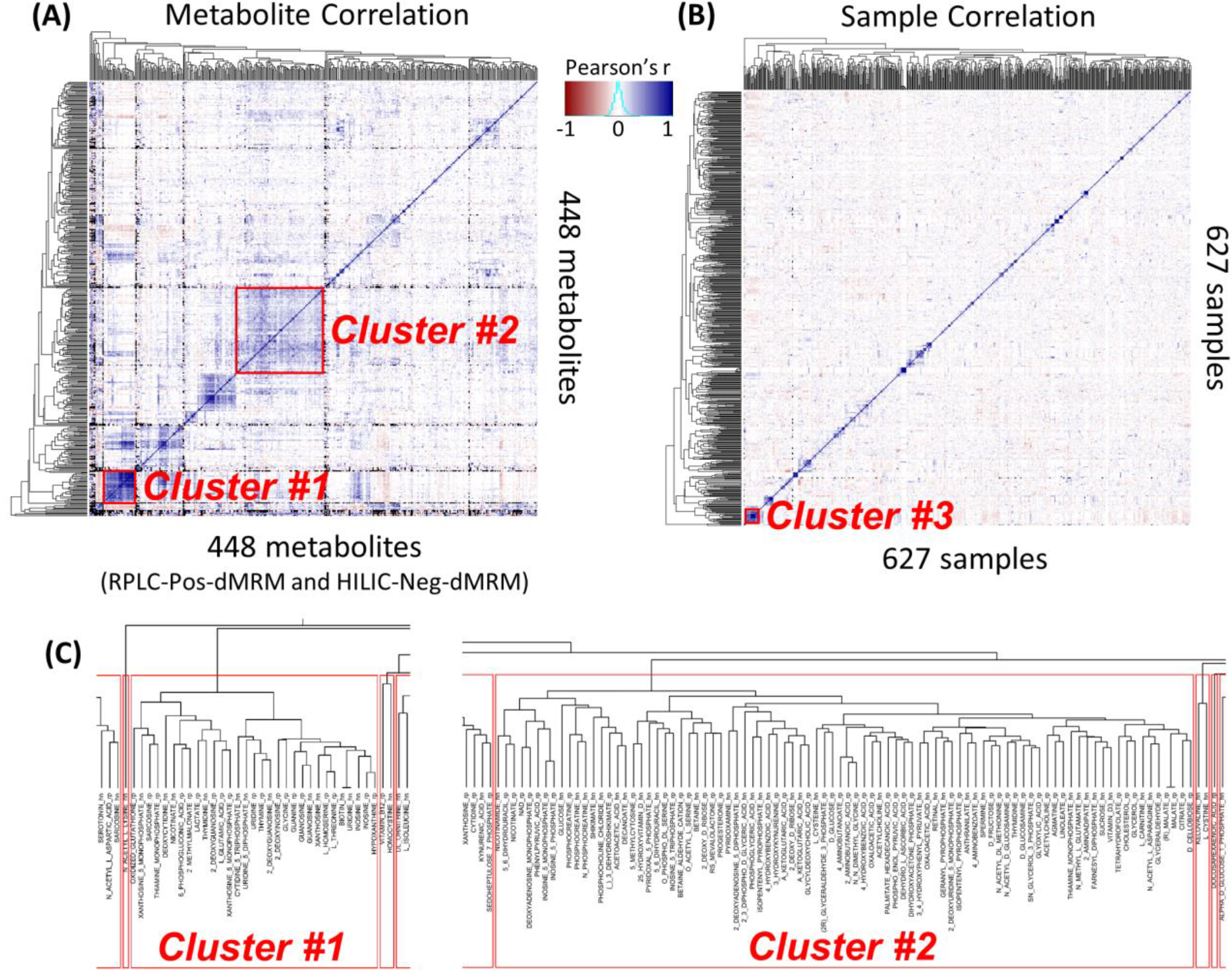
Global view of the metabolomics data compendium. (A) A correlation heatmap of 448 metabolite measurements of relative normalized abundance from both RPLC-Pos-dMRM and HILIC-Neg-dMRM methods along with unsupervised hierarchical clustering. The color key is based on Pearson correlation coefficients. (B) A correlation heatmap of 627 samples along with unsupervised hierarchical clustering. The color code is the same as in (A). See Supplementary Table S1 for the data used for (A) and (B). (C) Dendrogram trees of Cluster #1 and Cluster #2 at the height of 6. The suffix, “rp”, in the metabolite names stands for RPLC-Pos-dMRM and “hn” for HILIC-Neg-dMRM.

### 3.2. Analysis of normalized relative abundance

Raw data from the LC-MS analysis are peak areas of ion counts for each identified metabolite, which we use as relative abundance values. These raw data do not reflect absolute abundance and cannot be compared directly between experiments run at different times. Accordingly, we examine normalized relative abundance distributions of all metabolites across all datasets using medians in replicate-group measurements. Figure 2A shows the number of median measurements for each metabolite across all 183 replicate groups. There are a total of 13,026 measurements for 448 metabolites in both RPLC-Pos-dMRM and HILIC-Neg-dMRM methods. 71 metabolites were measured in all 183 replicate groups (Figure 2A). All of these came from the RPLC-Pos-dMRM method. This may indicate that the RPLC-Pos-dMRM method is more robust in detecting those metabolites than the HILIC-Neg-dMRM method. Those most frequently detected metabolites across different conditions are enriched in amino acids, nitrogen, glutathione, and purine metabolism (MetaboAnalyst; FDR < 0.01). The remaining metabolites show a monotonic decrease in the number of measured replicate groups. It is unclear what is the generating function or mechanism of this linearly decreasing distribution.

**Figure 2.**
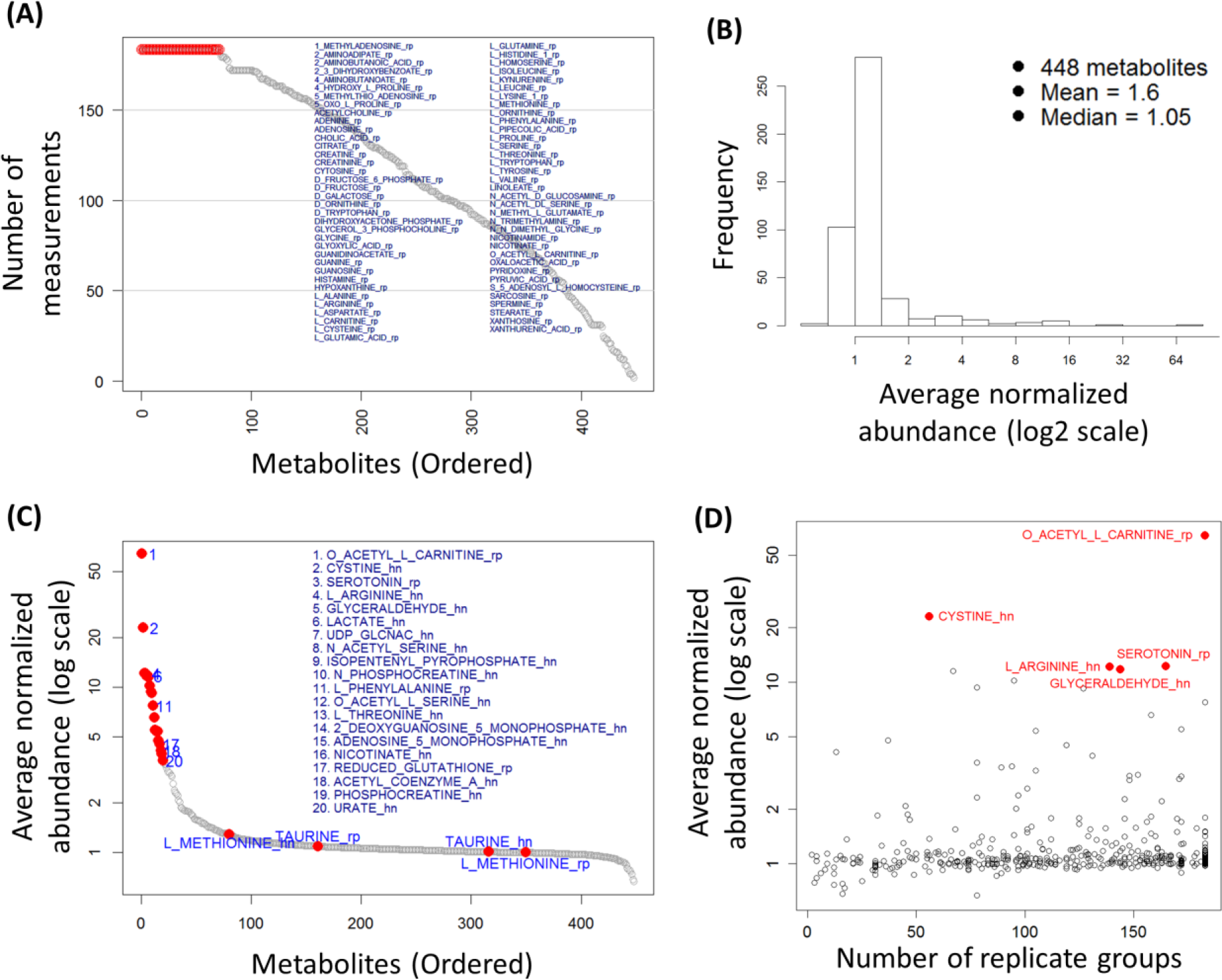
Analysis of relative abundance. (A) The distribution of the numbers of median measurements for 448 metabolites across all 183 biological replicate groups. There are 71 metabolites with measurements in the maximum 183 replicate groups as listed. See also Supplementary Table S2 for the list of 71 metabolites. (B) A histogram distribution of the average normalized abundance values for all 448 metabolites. (C) An ordered distribution of (B). The top 20 metabolites are listed along with methionine and taurine (cf. Figure 4F). See also Supplementary Table S2 for the list of the top 20 metabolites. (D) A scatter plot of the average normalized abundance of (B) or (C) and the numbers of replicate groups with measurements. The top 5 highly abundant metabolites are shown in red.

We next calculated the average of all abundance values for each metabolite. Figure 2B shows a histogram of the average abundance values for all 448 metabolites and Figure 2C shows a distribution of the ordered average abundance values. Three of the top 20 metabolites are involved in cysteine/methionine metabolism (cystine, acetyl-serine, and glutathione) and others in glycolysis, hexosamine, amino acids, mevalonate, energy, and redox metabolism. We also find that one of the top 20 metabolites, phenylalanine, is among the most reproducible metabolites by both RPLC-Pos-dMRM and HILIC-Neg-dMRM methods, along with methionine and taurine, as discussed below (Section 3.4). On the other hand, the bottom 20 metabolites include GDP-glucose, NADP+, fumaric acid, glycerol, dihydrofolate, leucine, docosahexaenoic acid, homocystine, TMP, sucrose, and deoxyadenosine. These metabolites are involved in many different metabolic pathways with no significant enrichment (MetaboAnalyst), suggesting that low abundance is not a pathway-specific or pathway-level feature. Figure 2D is a scatter plot of the average abundance and the number of replicate groups with measurements. The top 5 metabolites are highly abundant in 56 – 183 distinct groups on average, indicating the tendency of more frequent detection of more abundant metabolites. This analysis offers insight into the importance of those relatively high abundant metabolites in diverse conditions with potential effects on phenotypic differences.

### 3.3. Replicate-group CV analysis of RPLC-Pos-dMRM and HILIC-Neg-dMRM data

The coefficient of variation or CV (= *SD*/*mean*) is a standard quantitative measure for quality assessment in replicate groups. We calculated biological replicate-group CVs for both the RPLC-Pos-dMRM and HILIC-Neg-dMRM methods and examined their quality or reliability differences. Figure 3A shows a comparison of replicate-group CV distributions of the two methods. There are 28,765 CV values in RPLC-Pos-dMRM and 22,069 CV values in HILIC-Neg-dMRM from all 183 replicate groups for 220 and 228 metabolites, respectively. RPLC-Pos-dMRM measurements show more CVs < 0.2 than HILIC-Neg-dMRM measurements by about two-fold. The summary statistics also show overall better quality of RPLC-Pos-dMRM measurements than HILIC-Neg-dMRM. The reliability of each metabolite measurement was assessed by the average CV in all replicate groups. We denote the average CV of a metabolite by 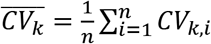 where *k* is a metabolite and *n* is the number of replicate groups with measurements. The ordered distributions of 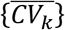 in Figure 3B shows an overview of average measurement reliability of all metabolites in each method (see also Supplementary Table S3). We calculated 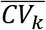 for only those metabolites that were measured in at least 20 replicate groups, which yielded 215 and 210 metabolites from RPLC-Pos-dMRM and HILIC-Neg-dMRM, respectively, with 154 of them in common. The reliability difference between the two methods is very clear, although Pearson’s correlation of CVs from the two methods for the 154 common metabolites shows a significant positive relationship (*r* = 0.37; *p* < 2.4e-6). This suggests greater chromatographic regularity or compound stability in RPLC-Pos-dMRM than HILIC-Neg-dMRM, which is consistent with previous studies on HILIC (Hao et al., 2008; Rhoades and Weljie, 2016). It also suggests that most metabolites show a similar tendency of reliability in each method. The quantity, 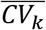, from our heterogeneous dataset could be used as a reliability index or a guidance in each method for other similar applications of LC-MS/MS targeted metabolomics. Among the 50 most reliable metabolites from each method, there are 20 overlapping metabolites including leucine/isoleucine, methionine, hydroxyproline, valine, nicotinamide, glutamate, phenylalanine, serine, glycine, and NAD (see Supplementary Table S3), which are likely to exhibit high stability and efficient ionizations in both methods. Among the 50 least reliable metabolites from each method, there are 14 overlapping metabolites including GDP-glucose, uridine 5’-diphosphate, cytidine 5’-diphosphate, sucrose, malate, guanosine 5’-diphosphate, lactate, and glyceraldehyde (see Supplementary Table S3), which would require more careful measurements and interpretations.

**Figure 3.**
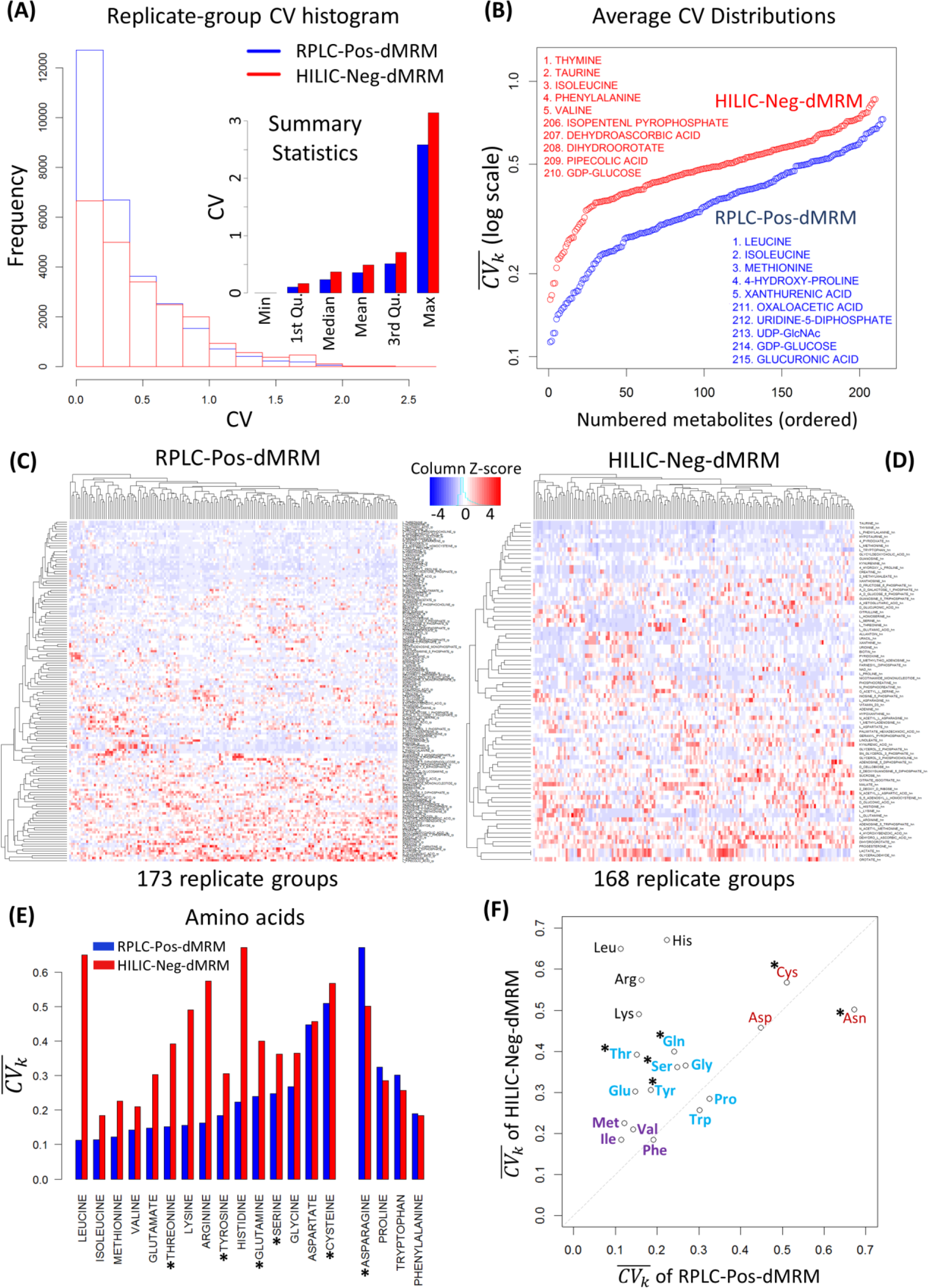
Replicate-group CV analysis. (A) Distributions of replicate-group CVs of the RPLC-Pos-dMRM and HILIC-Neg-dMRM methods. There are 28,765 CV values in RPLC-Pos-dMRM and 22,069 CV values in HILIC-Neg-dMRM from all 183 replicate groups for 220 and 228 metabolites, respectively. Summary statistics of all replicate-group CVs from the two methods are shown in the inset. (B) The ordered distributions of 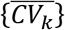 for individual metabolites from the RPLC-Pos-dMRM and HILIC-Neg-dMRM methods. The top 5 and bottom 5 metabolites at the two tails are listed from each method. (C) Heatmap and hierarchical clustering of replicate-group CVs for 145 metabolites with missing values less than 30% across all replicate groups in the RPLC-Pos-dMRM method. (D) Heatmap and hierarchical clustering of replicate-group CVs for 77 metabolites with missing values less than 30% across all replicate groups in the HILIC-Neg-dMRM method. CVs are standardized by the column Z-score in (C) and (D). (E) The bar graph shows 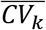 of amino acids. The 15 amino acids one the left possess lower 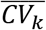 or better reproducibility in RPLC-Pos-dMRM and the 4 amino acids on the right possess lower 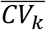 or better reproducibility in HILIC-Neg-dMRM. The six polar amino acids are indicated by asterisks. (F) A scatter plot of 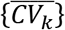 of the 19 amino acids in (E) shows a reproducibility trend and patterns as color coded as a guidance. The six polar amino acids are indicated by asterisks. See also Supplementary Table S3 for all 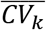 values in (B) and (E-F) and the ordered lists of the 145 and 77 metabolites of (C) and (D).

To examine the reliability in more detail, we further restricted our focus to those metabolites with CV missing values less than 30% for statistical robustness. There are 145 such metabolites for RPLC-Pos-dMRM and 77 such metabolites for HILIC-Neg-dMRM. The heatmaps and hierarchical clustering in Figures 3C and 3D show groups of metabolites with low CVs (i.e., better reliability; rows in blue) in each method. While we visually notice overall better reliability in RPLC-Pos-dMRM from the heatmaps, there are metabolites with low CVs across all replicate groups in HILIC measurements such as taurine, thymine, phenylalanine, hypotaurine, pyridoxate, methionine, and tryptophan.

A pathway analysis of most reliable metabolites with 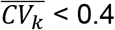 (125 and 54 metabolites in RPLC-Pos-dMRM and HILIC-Neg-dMRM, respectively) shows that each method does not favor any unique pathway. Both methods tend to have reliable measurements for amino acids and nucleotides biosynthesis pathways, the main difference being the number of reliable metabolites given a threshold of 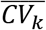. This pathway analysis supports the aforementioned positive correlation of 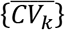 between the two methods. On the other hand, by focusing on 19 amino acids from both methods, we find that 4 amino acids show lower 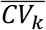 in HILIC-Neg-dMRM than in RPLC-Pos-dMRM: phenylalanine, tryptophan, proline, and asparagine, the first three of which show similarly good measurement reproducibility as hydrophobic in both methods with 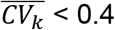 (Figures 3E and 3F). This suggests that those three amino acids may be used as most reliable reference metabolites in abundance measurements as well as in LC-MS method development in both chromatographic column conditions. It is also observed that asparagine, aspartate, and cysteine are less reliable in both methods (Figures 3E and 3F). We note that phenylalanine and tryptophan are often used as quality control standards in metabolomics laboratories on an empirical basis, corroborating our findings (Dr. Maureen Kachman, personal communication). We point out that all polar amino acids except asparagine show better reproducibility (i.e., lower 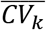) in RPLC-Pos-dMRM than HILIC-Neg-dMRM (Figures 3E and 3F).

### 3.4. Correlation analysis of RPLC-Pos-dMRM and HILIC-Neg-dMRM data

We continued to examine differences between RPLC-Pos-dMRM and HILIC-Neg-dMRM from a correlation point of view by focusing on those metabolites that were measured in at least 70% of all 42 experiments with both methods. We asked if abundance profiles of each metabolite from the two methods are well correlated in individual experiments as they were conducted on different days. A total of 3601 measurements were made for 237 metabolites in the 42 experiments. There were 221 metabolites that were measured in at least 2 experiments with both methods. 47 of them were measured in at least 70% of all 42 experiments. For each metabolite in each experiment, we calculated the Pearson correlation coefficient between two abundance profiles measured by the two methods. Figure 4A shows a heatmap of all RPLC-HILIC correlation coefficients, where the metabolites were sorted by the average correlation across all the experiments and the columns were sorted by the average correlation across all the metabolites. Larger circles in darker blues indicate good correlation between the two methods. The top metabolites with reliable measurements across most of the experiments include guanosine, 5-methlythio-adenosine, phosphocreatine, xanthine, proline, taurine, NAD+, and methionine, among others. On the other hand, linoleate, glucuronic acid, 4-hydroxy-L-proline, dehydro-L-ascorbic acid, and acetylcholine, among others show inconsistent measurements between the two methods, indicating that their retention mechanisms or ionization efficiency may greatly differ between the two methods from experiment to experiment on different days.

**Figure 4.**
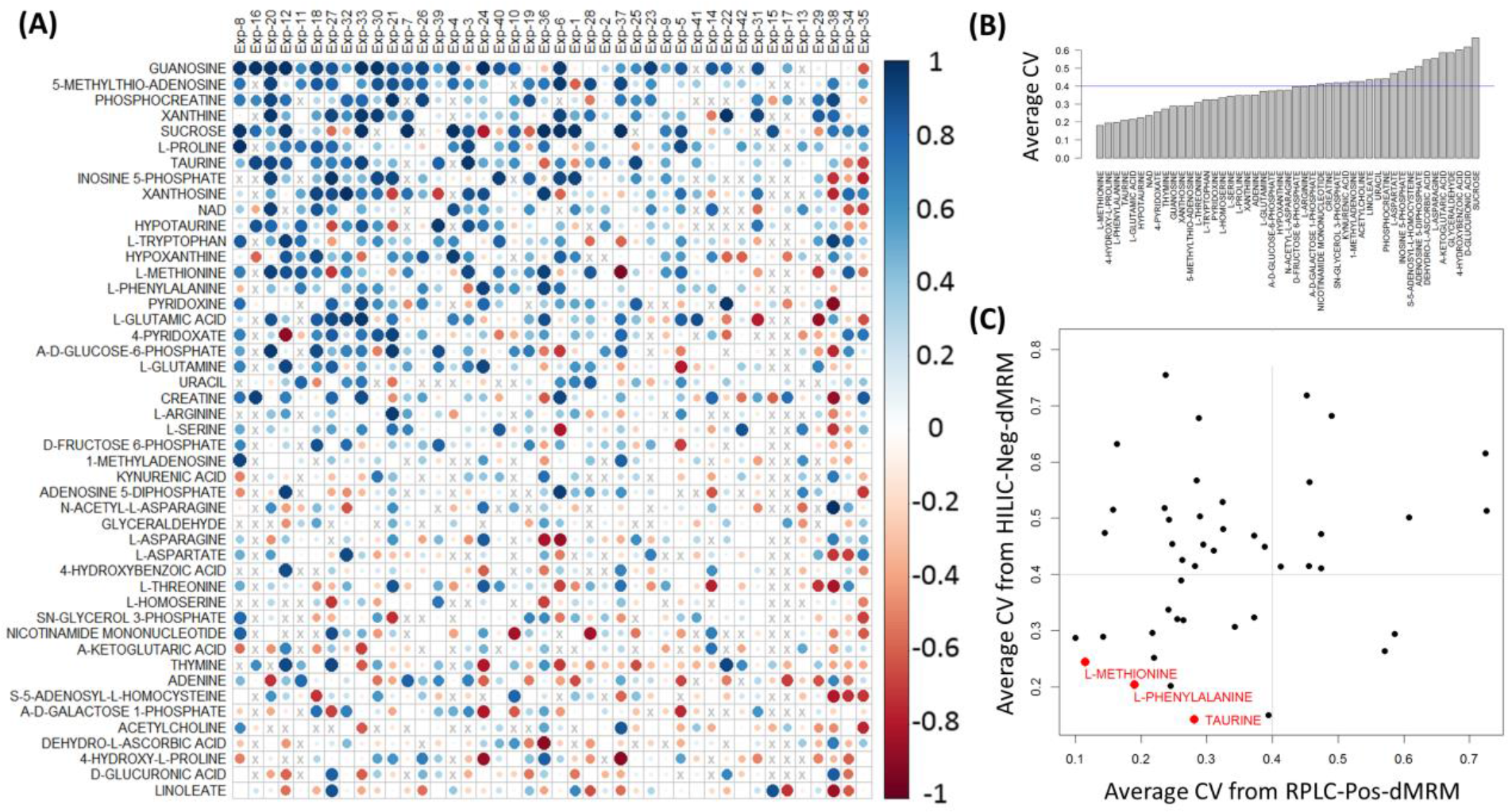
RPLC-HILIC correlation analysis. (A) Heatmap of all RPLC-HILIC Pearson correlation coefficients for 47 metabolites that were measured in at least 70% of all 42 experiments. The metabolites were sorted by the average correlation across all the experiments and the columns were sorted by the average correlation across all the metabolites. Larger circles in darker blues indicate good correlation between the RPLC-Pos-dMRM and HILIC-Neg-dMRM measurements. (B) Distribution of 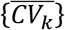 across the 42 datasets in both RPLC-Pos-dMRM and HILIC-Neg-dMRM methods. (C) A scatter plot of 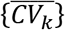 from the RPLC-Pos-dMRM and HILIC-Neg-dMRM measurements for all 47 metabolites. The 3 most reproducible metabolites by both methods are shown in red. See also Supplementary Table S4 for the list of 47 metabolites along with 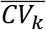 in both methods.

To further examine the 47 metabolites measured by the two methods, we next analyzed 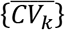 as in the previous section. Figure 4B shows a distribution of 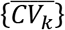, where those metabolites on the left with smaller 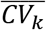 are more reproducible and hence reliable across the 42 datasets by both methods on average. The top 7 metabolites are methionine, 4-hydroxy-L-proline, phenylalanine, taurine, glutamic acid, hypotaurine, and NAD+ with 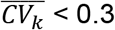. We also calculated the average of 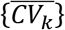 in each of the two methods. The scatter plot in Figure 4C shows a distribution of 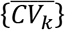 from the two methods for all 47 metabolites. Given our reference value of 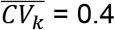, there are 15 metabolites with 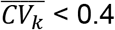 in both methods that we deem reproducible across diverse conditions. The 3 most reproducible metabolites from both methods are methionine, phenylalanine, and taurine. Phenylalanine was among the top 20 metabolites of high average abundance as discussed above. There are several metabolites which are reliable in either method alone based on a threshold of 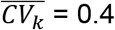. The RPLC-Pos-dMRM measurements are more reliable for 35 metabolites, whereas the HILIC-Neg-dMRM measurements are more reliable for 17 metabolites.

### 3.5. Analysis of abundance fold changes for effect size and variability

To study the directionality and magnitude of metabolite changes upon experimental perturbation, we analyzed fold changes (FC) of metabolite abundance across all case-control condition group pairs. For all case-control pairs and each metabolite, we calculated the median 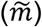 of normalized abundance values in each group of replicates, i.e.,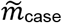 and 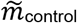, and the fold change 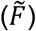 of the medians, 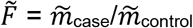. Then, we calculated the average of absolute values of 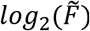 for all case-control pairs for each metabolite, 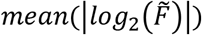, which represents an average magnitude of the fold change in the case group relative to the control group for that metabolite. The magnitude or absolute value of 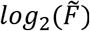 is also called the effect size.

There are a total of 448 metabolites in 176 case-control pairs from both RPLC-Pos-dMRM and HILIC-Neg-dMRM methods. There are 37.3% missing values in the 448 by 176 data matrix. A missing value occurs when no measurement was made in either a case or control condition. For a hierarchical clustering of the full data, we removed those metabolites and case-control pairs that have more than 50% missing values across all case-control pairs and all metabolites, respectively. This left us with 294 metabolites and 161 case-control pairs with 15.8% missing values. Figure 5A shows a heatmap of the 294 by 161 data matrix of 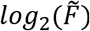 values along with hierarchical clustering dendrograms on both axes. The full 448 by 176 data matrix of fold changes gives us a histogram of all average fold-change magnitudes defined by 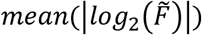 and their ordered distribution in Figure 5B. The average fold-change magnitudes for most metabolites are less than 2 (the peak of the histogram) and the top 20 metabolites with the highest average effect sizes are mostly involved in glycolysis, redox, pyrimidine, and methionine/cysteine/folate metabolism. Homocystine shows the largest average FC magnitude of more than 20, and lactate/glyceraldehyde the second largest of more than 9. We note that lactate and glyceraldehyde were not distinguishable in our HILIC-Neg-dMRM method with identical data. Most of the top 20 metabolites are measured in more than 70 case-control pairs, except homocystine in 9 pairs in the HILIC method and NADP in 12 pairs in the HILIC method (Figure 5C). In addition, we calculated the standard deviation (SD) of the effect sizes for each metabolite to examine the effect-size variability in all case-control pairs for that metabolite. Figure 5D shows a histogram of the SDs and an ordered SD distribution. The SDs for most metabolites are less than 1 and there are 7 metabolites with SD > 3 including lactate/glyceraldehyde and orotate. We also performed an analysis of median absolute deviation (MAD) to complement the SD analysis. Figure 5E shows a MAD histogram and an ordered MAD distribution. The top 20 most variable metabolites with the highest MADs are involved in glycolysis, redox, pyrimidine, energy, and methionine/cysteine/folate metabolism. Figures 5F and 5G show good positive correlations between the average effect sizes and the SD and MAD variability, respectively. Homocystine, lactate/glyceraldehyde, orotate, GSH, and dihydrofolate have both the highest average effect size and the largest variability by both SD and MAD among the top 20, which suggests that they are most dynamic and responsive metabolites across this diverse array of treatments.

**Figure 5.**
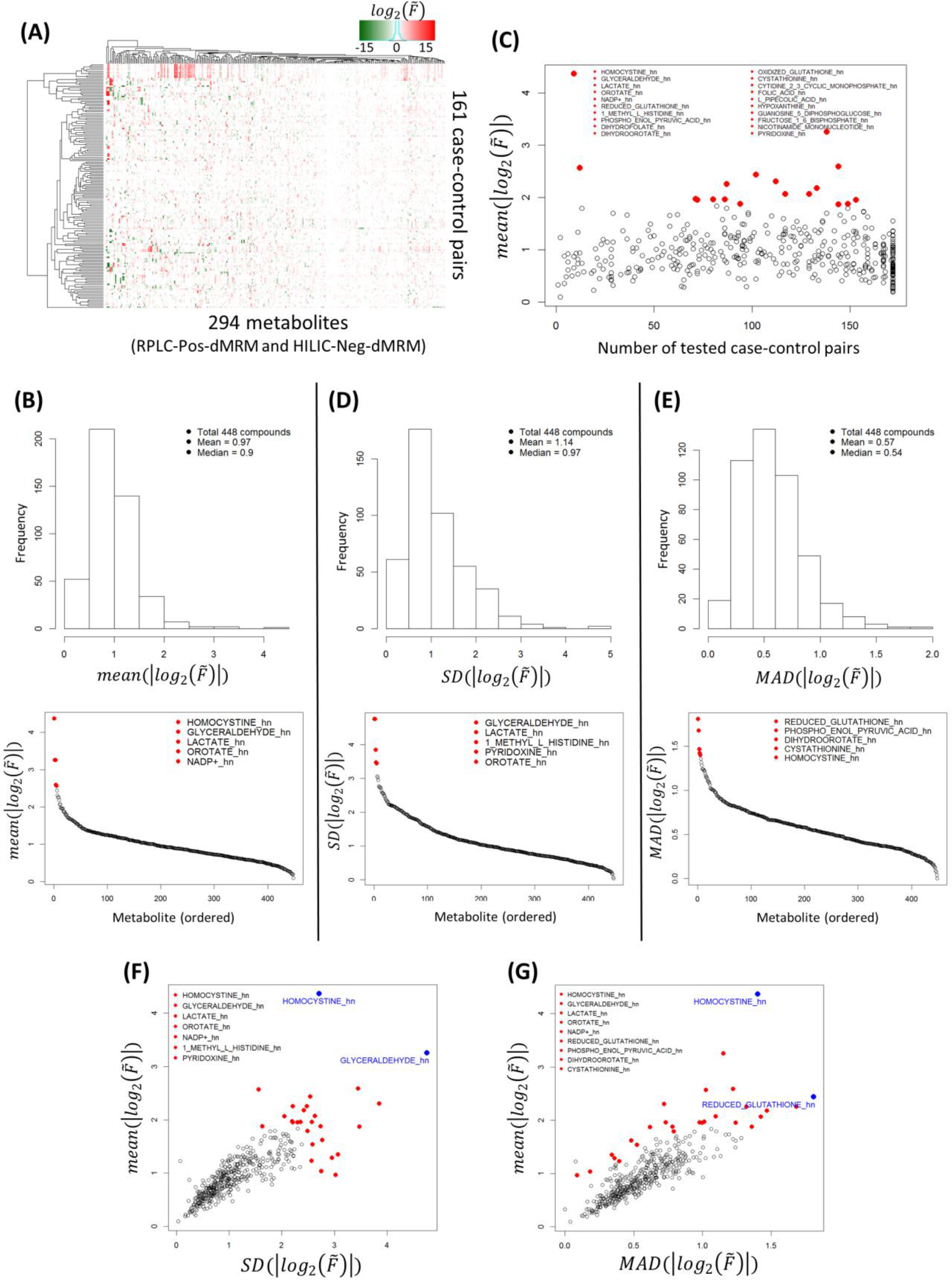
Abundance fold change analysis. (A) Heatmap of 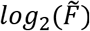 for 294 metabolites and 161 case-control sample pairs. (B) A histogram of all average fold-change magnitudes (=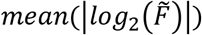; effect size) and an ordered distribution of the average magnitudes. The top 20 metabolites are listed. (C) A scatter plot of the average fold-change magnitudes and the numbers of tested case-control pairs. (D) Effect-size variability in terms of the standard deviation (SD). A histogram of SDs of the effect sizes and an ordered SD distribution. The top 5 metabolites are listed. (E) Effect-size variability in terms of the maximum absolute deviation (MAD). A histogram of MADs of the effect sizes and an ordered MAD distribution. The top 5 metabolites are listed. (F) A scatter plot of the average effect sizes and the SD variability. The union of the top 5 metabolites from (B) and (D) are listed. (G) A scatter plot of the average effect sizes and the MAD variability. The union of the top 5 metabolites from (B) and (E) are listed. See Supplementary Table S5 for the full fold-change dataset.

## 4. Conclusions

We performed LC-MS/MS targeted metabolomics to generate 42 datasets in a wide range of experiments and sample types from the same platform in both RPLC-Pos-dMRM and HILIC-Neg-dMRM methods. We carried out a series of computational and statistical analyses to systematically assess data quality of both methods in biological replicate groups to identify more reliable measurements in each method. We find that the RPLC-Pos-dMRM method tends to generate more reproducible measurements than the HILIC-Neg-dMRM method, including polar amino acids. On the other hand, several metabolites are measured reliably in both methods such as methionine, phenylalanine, and taurine. Phenylalanine is among the most abundant metabolites across all samples on average along with acetylcarnitine, cystine, arginine, lactate/glyceraldehyde, UDP-GlcNAc, AMP, GSH, and acetyl-CoA. We also characterized global abundance changes and variability in case-control groups to identify most dynamic and variable metabolites across heterogeneous conditions, such as homocystine, lactate/glyceraldehyde, and GSH. Those metabolites are involved in cysteine metabolism (taurine, cystine, homocystine), glycolysis (glyceraldehyde, lactate), hexosamine biosynthesis (UDP-GlcNAc), redox balance (GSH), fatty acids metabolism (acetylcarnitine, acetyl-CoA), and the TCA cycle (acetyl-CoA). This might suggest that they tend to play more significant roles than others in the associated metabolic pathways under many different conditions. For example, lactate, taurine, methionine, and acetylcarnitine were recently found to be more frequently differentially abundant in tumors compared to normal tissues across different cancer types (Reznik et al., 2018). Our study offers a systematic guideline and a reference point in targeted metabolomics analysis by either RPLC-Pos-dMRM or HILIC-Neg-dMRM method for more reliable biological interpretations without sample-dependent optimization of the LC-MS system.

## Supplementary material

A list of metabolites measured in RPLC-Pos-dMRM and HILIC-Neg-dMRM methods is available in the Supplementary Table along with their formulae, precursor ion mass, product ion mass, RTs, RT window, and CAS numbers. The compendium of randomized and deidentified abundance data is available in Supplementary Table S1, which was used in Figures 1A and 1B. The fold-change dataset of case-control condition pairs used in Figure 5 is available in Supplementary Table S5 after randomization and deidentification of the condition pairs. All analysis codes in R are available upon request.

## Acknowledgements

The authors would like to thank all of anonymous collaborators who provided the samples used in this study and Drs. Garrison Birch, Steven Fischer, and Maureen Kachman for their critical comments and suggestions. D.M.K is supported by the University of Michigan's Program in Chemical Biology Graduate Assistance in Areas of National Need (GAANN) award. C.A.L. was supported by a Pancreatic Cancer Action Network/AACR Pathway to Leadership award (13-70-25-LYSS); Dale F. Frey Award for Breakthrough Scientists from the Damon Runyon Cancer Research Foundation (DFS-09-14); Junior Scholar Award from The V Foundation for Cancer Research (V2016-009); Kimmel Scholar Award from the Sidney Kimmel Foundation for Cancer Research (SKF-16-005); a 2017 AACR NextGen Grant for Transformative Cancer Research (17-20-01-LYSS); and UMCCC Core Grant (P30 CA046592). Metabolomics studies performed at the University of Michigan were supported by NIH grant DK097153, the Charles Woodson Research Fund and the UM Pediatric Brain Tumor Initiative.

## Author contributions

HL conceived the idea, designed the algorithms and pipelines, performed computational and statistical analyses, interpreted the results, and wrote a draft of the manuscript. DMK and PS prepared the samples and helped with raw data processing. LZ performed LC-MS analysis including raw data generation. DMK and LZ contributed to writing the manuscript. CAL conceived the idea, interpreted the results, contributed to writing the manuscript, and supervised the project. All authors read and approved the manuscript.

## Compliance with ethical standards

This article does not contain any studies with human and/or animal participants performed by any of the authors.

## References

Buszewski, B., and Noga, S. (2012). Hydrophilic interaction liquid chromatography (HILIC)”-a powerful separation technique. Anal Bioanal Chem 402, 231–247.

Chong, J., Soufan, O., Li, C., Caraus, I., Li, S., Bourque, G., Wishart, D.S., and Xia, J. (2018). MetaboAnalyst 4.0: towards more transparent and integrative metabolomics analysis. Nucleic Acids Research 46, W486–W494.

Cubbon, S., Antonio, C., Wilson, J., and Thomas-Oates J. (2010). Metabolomic applications of HILIC-LC-MS. Mass Spectrom Rev 29, 671–684.

Dudley, E., Yousef, M., Wang, Y., and Griffiths, W.J. (2010). Targeted metabolomics and mass spectrometry. Adv Protein Chem Struct Biol 80, 45–83.

Fuhrer, T., and Zamboni, N. (2015). High-throughput discovery metabolomics. Curr Opin Biotechnol 31, 73–78.

Griffiths, W.J., Koal, T., Wang, Y., Kohl, M., Enot, D.P., and Deigner, H.P. (2010). Targeted metabolomics for biomarker discovery. Angew Chem Int Ed Engl 49, 5426–5445.

Halbrook, C.J., Pontious, C., Lee, H.-J., Kovalenko, I., Zhang, Y., Lapienyte, L., Dreyer, S., Kremer, D.M., Sajjakulnukit, P., Zhang, L., et al. (2018). Macrophage Released Pyrimidines Inhibit Gemcitabine Therapy in Pancreatic Cancer. bioRxiv, 463125.

Hao, Z., Xiao, B., and Weng, N. (2008). Impact of column temperature and mobile phase components on selectivity of hydrophilic interaction chromatography (HILIC). Journal of separation science 31, 1449–1464.

Ivanisevic, J., Zhu, Z.J., Plate, L., Tautenhahn, R., Chen, S., O’Brien, P.J., Johnson, C.H., Marletta, M.A., Patti, G.J., and Siuzdak, G. (2013). Toward ‘omic scale metabolite profiling: a dual separation-mass spectrometry approach for coverage of lipid and central carbon metabolism. Anal Chem 85, 6876–6884.

Johnson, C.H., Ivanisevic, J., and Siuzdak, G. (2016). Metabolomics: beyond biomarkers and towards mechanisms. Nat Rev Mol Cell Biol 17, 451–459.

Lu, W., Bennett, B.D., and Rabinowitz, J.D. (2008). Analytical strategies for LC-MS-based targeted metabolomics. J Chromatogr B Analyt Technol Biomed Life Sci 871, 236–242.

Patti, G.J., Yanes, O., and Siuzdak, G. (2012). Innovation: Metabolomics: the apogee of the omics trilogy. Nat Rev Mol Cell Biol 13, 263–269.

Reznik, E., Luna, A., Aksoy, B.A., Liu, E.M., La, K., Ostrovnaya, I., Creighton, C.J., Hakimi, A.A., and Sander, C. (2018). A Landscape of Metabolic Variation across Tumor Types. Cell Syst 6, 301–313.e303.

Rhoades, S.D., and Weljie, A.M. (2016). Comprehensive Optimization of LC-MS Metabolomics Methods Using Design of Experiments (COLMeD). Metabolomics : Official journal of the Metabolomic Society 12.

Rojo, D., Barbas, C., and Ruperez, F.J. (2012). LC-MS metabolomics of polar compounds. Bioanalysis 4, 1235–1243.

Schofield, H.K., Zeller, J., Espinoza, C., Halbrook, C.J., Del Vecchio, A., Magnuson, B., Fabo, T., Daylan, A.E.C., Kovalenko, I., Lee, H.J., et al. (2018). Mutant p53R270H drives altered metabolism and increased invasion in pancreatic ductal adenocarcinoma. JCI Insight 3.

Sousa, C.M., Biancur, D.E., Wang, X., Halbrook, C.J., Sherman, M.H., Zhang, L., Kremer, D., Hwang, R.F., Witkiewicz, A.K., Ying, H., et al. (2016). Pancreatic stellate cells support tumour metabolism through autophagic alanine secretion. Nature 536, 479–483.

Svoboda, L.K., Teh, S.S.K., Sud, S., Kerk, S., Zebolsky, A., Treichel, S., Thomas, D., Halbrook, C.J., Lee, H.J., Kremer, D., et al. (2018). Menin regulates the serine biosynthetic pathway in Ewing sarcoma. J Pathol 245, 324–336.

Tang, D.Q., Zou, L., Yin, X.X., and Ong, C.N. (2016). HILIC-MS for metabolomics: An attractive and complementary approach to RPLC-MS. Mass Spectrom Rev 35, 574–600.

Wishart, D.S. (2016). Emerging applications of metabolomics in drug discovery and precision medicine. Nat Rev Drug Discov 15, 473–484.

Xie, B., Wang, Y., Jones, D.R., Dey, K.K., Wang, X., Li, Y., Cho, J.H., Shaw, T.I., Tan, H., and Peng, J. (2018). Isotope Labeling-Assisted Evaluation of Hydrophilic and Hydrophobic Liquid Chromatograph-Mass Spectrometry for Metabolomics Profiling. Anal Chem 90, 8538–8545.

Yuan, M., Breitkopf, S.B., Yang, X., and Asara, J.M. (2012). A positive/negative ion-switching, targeted mass spectrometry-based metabolomics platform for bodily fluids, cells, and fresh and fixed tissue. Nat Protoc 7, 872–881.

Zhou, J., and Yin, Y. (2016). Strategies for large-scale targeted metabolomics quantification by liquid chromatography-mass spectrometry. Analyst 141, 6362–6373.

